# Generation of a human *Tropomyosin 1* knockout iPSC line

**DOI:** 10.1101/2023.05.03.539242

**Authors:** Madison B Wilken, Jean Ann Maguire, Lea V Dungan, Alyssa Gagne, Catherine Osorio-Quintero, Elisa A Waxman, Stella T Chou, Paul Gadue, Deborah L French, Christopher S Thom

## Abstract

The CHOPWT17_TPM1KOc28 iPSC line was generated to interrogate the functions of *Tropomyosin 1* (*TPM1*) in primary human cell development. This line was reprogrammed from a previously published wild type control iPSC line.

## Resource Table

|  |  |
| --- | --- |
| Unique stem cell line identifier | CHOPi013-A |
| Alternate name of stem cell line | CHOPWT17_TPM1KOc28 |
| Institution | The Children's Hospital of Philadelphia (CHOP),<br>Philadelphia, Pennsylvania, USA |
| Contact information of distributor | Christopher S. Thom, |
| Type of cell line | iPSC |
| Origin | Human |
| Additional origin info | Age: 51 years Sex: Male Ethnicity: Caucasian Type: O- |
| Cell source | Blood |
| Clonality | Clonal |
| Method of reprogramming | N/A |
| Genetic modification | CRISPR-Cas9 |
| Associated disease | N/A |
| Gene/Locus | TPM1 |
| Method of modification | N/A |
| Name of transgene or resistance | N/A |
| Inducible/constitutive system | N/A |
| Date archived/stock date | February 2022 |
| Cell line repository/bank | https://hpscreg.eu/cell-line/CHOPi013-A |
| Ethical approval | Source cells purchased from BioIVT (Cell Line #1000,<br>HUMANWBNE1803609) |

## Resource Utility

The CHOPWT17_TPM1KOc28 iPSC line was generated to interrogate *Tropomyosin 1* (*TPM1*) biology (Gunning and Hardeman, 2017). The actin-regulatory *TPM1* gene impacts blood traits and developmental hematopoiesis (Thom et al., 2020). This iPSC line can facilitate study of *TPM1* in multiple developmental contexts.

## Resource Details

Whole blood (BioIVT, Cell Line #1000, HUMANWBNE1803609) from a 51-year-old Caucasian male was used to generate a previously reported parental CHOPWT17 iPSC line (An et al., 2022). Using CRISPR/CAS9 genome editing of CHOPWT17 iPSCs, a homozygous deletion at codon 304 (deleted C) was introduced and confirmed by DNA sequencing (**Fig. 1A**). This resulted in a frameshift (fs) mutation that caused a premature stop codon at position 127 (X127). Western blot confirmed *TPM1* knockout (**Fig. 1B**). Thus, this iPSC line abrogates *TPM1* expression in iPSCs without loss of intronic regulatory regions, which differs from previously published iPSC lines (Thom et al., 2020). This selected clone, CHOPWT17_TPM1KOc28, was then expanded, cryopreserved at passage 40, and subsequently analyzed (Tables 1 and 2). CHOPWT17_TPM1KOc28 iPSCs showed cellular morphology and intranuclear SOX2 expression identical to its wild-type parental iPSC line (**Fig. 1C and 1D**). Expression of stem cell surface markers was normal by flow cytometry (**Fig. 1E**). A normal karyotype (46, XY) was revealed by G-band analysis (**Fig. 1F**). Pluripotency potential was shown by differentiation into three germ layers, followed by flow cytometric analyses that confirmed relevant cell surface marker expression (**Fig. 1G** and **Supp Fig. 1A**). DNA fingerprinting by STR analysis authenticated the genetic identity of the iPSC line in relation the parental cell line (**Supp Fig. 1B**) and CHOPWT17_TPM1KOc28 iPSCs tested negative for Mycoplasma (**Supp Fig. 1C**).

**Table 1.** Characterization and validation.

| **Classification** | **Test** | **Result** | **Data** |
| --- | --- | --- | --- |
| Mutation Analysis | Sequencing | Homozygous c.304delG (p.A102PfsX127) | Fig. 1A |
|  | Western blot | Absent TPM1 | Fig. 1B |
| Morphology | Photography | Normal | Fig. 1C |
| Phenotype | Immunocytochemistry | Staining of marker SOX2 | Fig. 1D |
|  | Flow cytometry | SEA3/4, Tra1–60/81 | Fig. 1E |
| Genotype | Karyotype (G-Banding) and resolution | Normal, XY; Resolution: fair | Fig. 1F |
| Differentiation potential | *In vitro* differentiation | FACS staining for KDR/CD31 (mesoderm/endothelium), CD117/CD184 (endoderm), or SOX2/PAX6 (ectoderm). FOXG1 also confirmed ectoderm identity | Fig. 1G and Supp Fig. 1A |
| Identity | STR Analysis | Matched iPSCs to parental line | Supp Fig. 1B |
| Microbiology and virology | Mycoplasma | Negative by PCR | Supp. Fig. 1C |
| Donor screening | HIV 1 + 2, Hepatitis B, Hepatitis C | N/A | N/A |
| Genotype additional info | Blood group genotyping | N/A | N/A |
|  | HLA tissue typing | N/A | N/A |

**Table 2:**
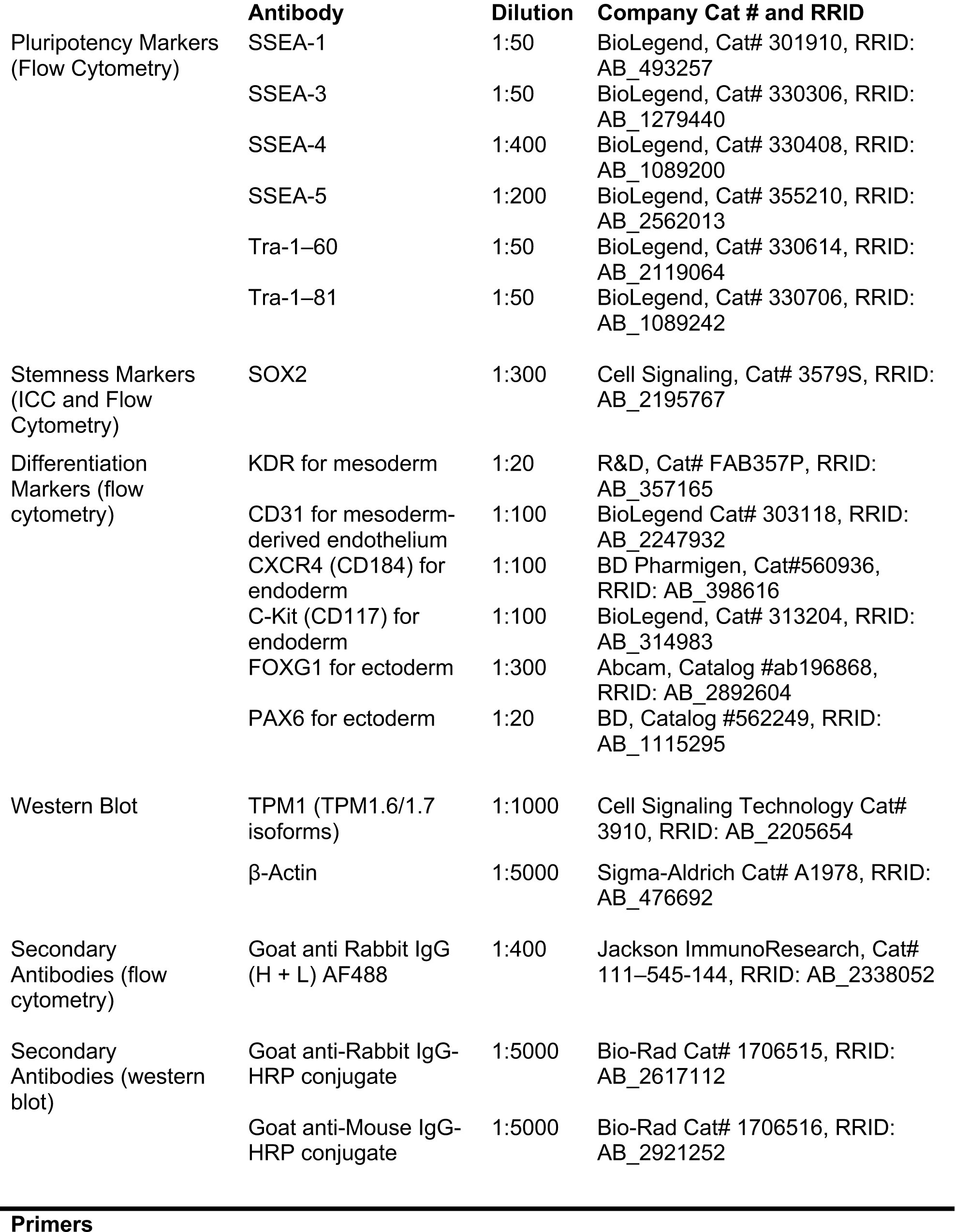
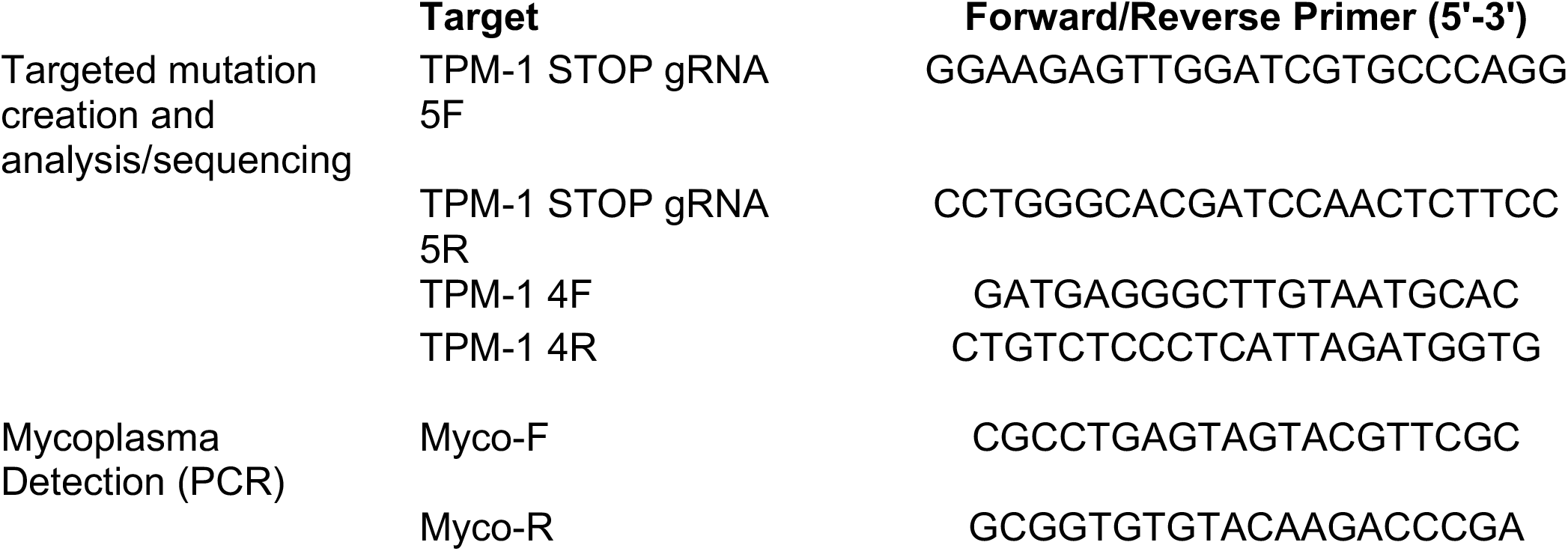
Reagent details.

**Figure 1.**
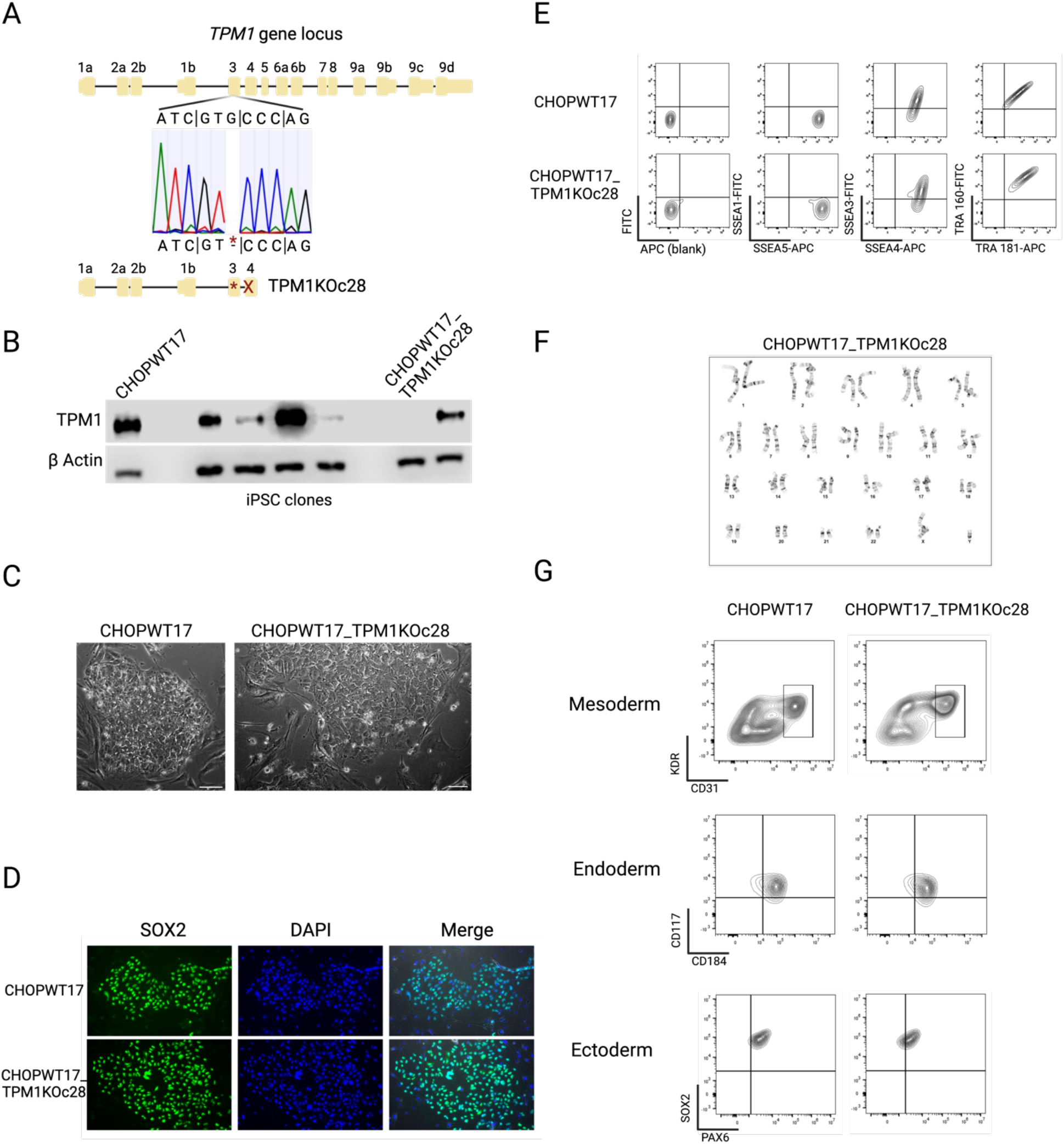
Characterization of the human iPSC line CHOPWT17_TPM1KOc28. A. Sequencing analysis of the targeted iPSC clone showing homozygous deletion at the indicated site. B. Western blot showing absence of TPM1 protein in the selected clone. C. Morphologic appearance of CHOPWT17_TPM1KOc28 iPSC clones match the parental cell line. Scale bar, 50 μm. D. Immunofluorescence staining confirms intranuclear SOX2 staining in CHOPWT17_TPM1KOc28 iPSCs and parental cells. Scale bar, 50 μm. E. Flow cytometry analysis shows appropriate stem cell marker expression in CHOPWT17_TPM1KOc28 iPSCs and parental cells. F. Representative normal karyotype for CHOPWT17_TPM1KOc28 iPSCs. G. Directed differentiation shows efficient production of mesoderm (including mesoderm-derived KDR^+^/CD31^+^ endothelial cells shown in box), endoderm, and ectoderm.

## Materials and Methods

1. **Cell Culture** The iPSCs were maintained on irradiated Mouse Embryonic Fibroblasts (MEFs) in 6-well tissue culture plates at 37^°^C with 5% CO_2_ and 5% O_2_ They were grown in human embryonic stem cell (hESC-10) medium made up of a base of DMEM/F12 (50:50 Gibco) with 20% knock-out serum replacement (KOSR, Gibco), 100 μM nonessential amino acids, 2 mM glutamine, 50 U/ml penicillin, 50 μg/ml streptomycin (all from Invitrogen), 10^−4^ M β–mercaptoethanol (Sigma, St. Louis, MO), and 10 ng/ml human bFGF (BioTechne). Cells were passaged every 5-7 days once they reached about 70-80% confluency using TrypLE cell dissociation reagent (Gibco) and gentle scraping before being replated onto new MEFs. Cells were transitioned in hESC-10 medium containing with 10 μM ROCK inhibitor (Y27632 dihydrochloride, Tocris) for <24 hours before replacing the media with fresh hESC-10 medium without ROCK inhibitor. Directed differentiations were performed essentially as described for mesoderm (Thom et al., 2020), endoderm (Leavens et al., 2021), and ectoderm (Telezhkin et al., 2016).
2. **Primary cell source and CRISPR/CAS9 genome editing** The reprogrammed parental iPSC line (CHOPWT17.6) has been previously reported (An et al., 2022). Using a CRISPR/CAS9 protocol (Maguire et al., 2022), guide RNAs were designed to target exon 3 of the *TPM1* gene (GGAAGAGTTGGATCGTGCCCAGG, CCTGGGCACGATCCAACTCTTCC). The parental cell line was plated on irradiated MEFs to reach 80-85% confluency after an overnight incubation. The following quantities were used for transfection: 50 μl base medium (DMEM/F12), 0.5 μg Cas9-GFP, 0.5 μg of each gRNA and 3 μl Lipofectamine Stem Reagent. After 48 hours, FACS-sorted GFP^+^ cells were plated on a 10cm plate containing 1:3 matrigel and MEFs, then allowed to grow until colonies were established. Colonies were manually picked and expanded for screening. Targeted clones were identified using PCR-based screening strategies designed to detect editing. The chosen clone that suggested homozygous knockout was then sequenced and further analyzed to confirm *TPM1* knockout.
3. **Karyotype analysis** Cytogenic analysis was performed on twenty G-banded metaphase cells by Cell Line Genetics, Inc (Madison, WI). All 20 cells demonstrated a normal male karyotype.
4. **Flow cytometry** Single cells, collected using trypsin dissociation, were stained and analyzed with a CytoFLEX flow cytometer (Beckman Coulter) and FlowJo software (BD). Cells were incubated in the dark for 15 min at room temperature with the following antibodies to evaluate pluripotency expression: Alexa-Fluor®-647 α-human SSEA4 (1:400) and Alexa-Fluor®-488 SSEA3 (1:50); Tra-1–81 (1:50) and Tra-1–60 (1:50); Alexa-Fluor®-488 SSEA1 (CD15) (1:50) and Alexa-Fluor®-647 SSEA5 (1:200) (BioLegend). In all experiments, an unstained sample was used as a negative control.
5. **Western blot** Cell pellets from selected iPSC clones were lysed in RIPA buffer (Cell Signaling Technologies) and proteins electrophoretically separated on Nu-PAGE polyacrylamide gels (Invitrogen). Proteins were transferred to nitrocellulose membranes (Invitrogen), blocked in 3% milk in TBS-Tween, and probed with antibodies to detect TPM1 (1:1000, Cell Signaling Technologies) or β-Actin (1:5000, Sigma). Secondary antibodies were anti-Rabbit IgG-HRP or anti-Mouse IgG-HRP (Bio-Rad). Western blots were imaged using a Chemi-Doc Imaging System (Bio-Rad).
6. **Immunocytochemistry** Cells were plated in hESC-10 media on irradiated MEF-coated coverslips at 90% confluency and fixed in 4% paraformaldehyde for 15 min at room temperature. Permeabilization was performed using ice cold 100% methanol for 10 min at −20°C. Blocking and antibody buffers were prepared in 0.3% Triton-X100/PBS with 5% goat serum and Hank’s Balanced Salt Solution (HBSS), respectively. Cells were blocked at room temperature for 1 h with gentle rocking. Primary antibody (SOX2) incubation was performed at 4°C overnight, and secondary antibody (Goat α-Rabbit AF488) incubation at room temperature for 2 h. Coverslips were mounted in Vectashield mounting media with DAPI (Vector Laboratories, Burlingame, CA). Staining was visualized on an Olympus IX70 microscope and imaged with Metamorph (Molecular Devices).
7. **STR analysis** The genetic integrity of the iPSC line was confirmed by DNA fingerprinting. STR analysis was done by Cell Line Genetics, Inc (Madison, WI).
8. **Mycoplasma testing** Mycoplasma testing was performed by PCR amplification on iPSC genomic DNA. Cycling parameters were as follows: 95°C for 10 min, 35 cycles of 95°C 45 sec, 55°C 30 sec, 72°C 30 sec, a 72°C 10-minute extension, and a 4°C hold. PCR products were separated on a 1.0% agarose gel and visualized with ethidium bromide. Positive, and negative controls were used.

## Figure and Supplementary Figure

**Supplementary Figure 1.**
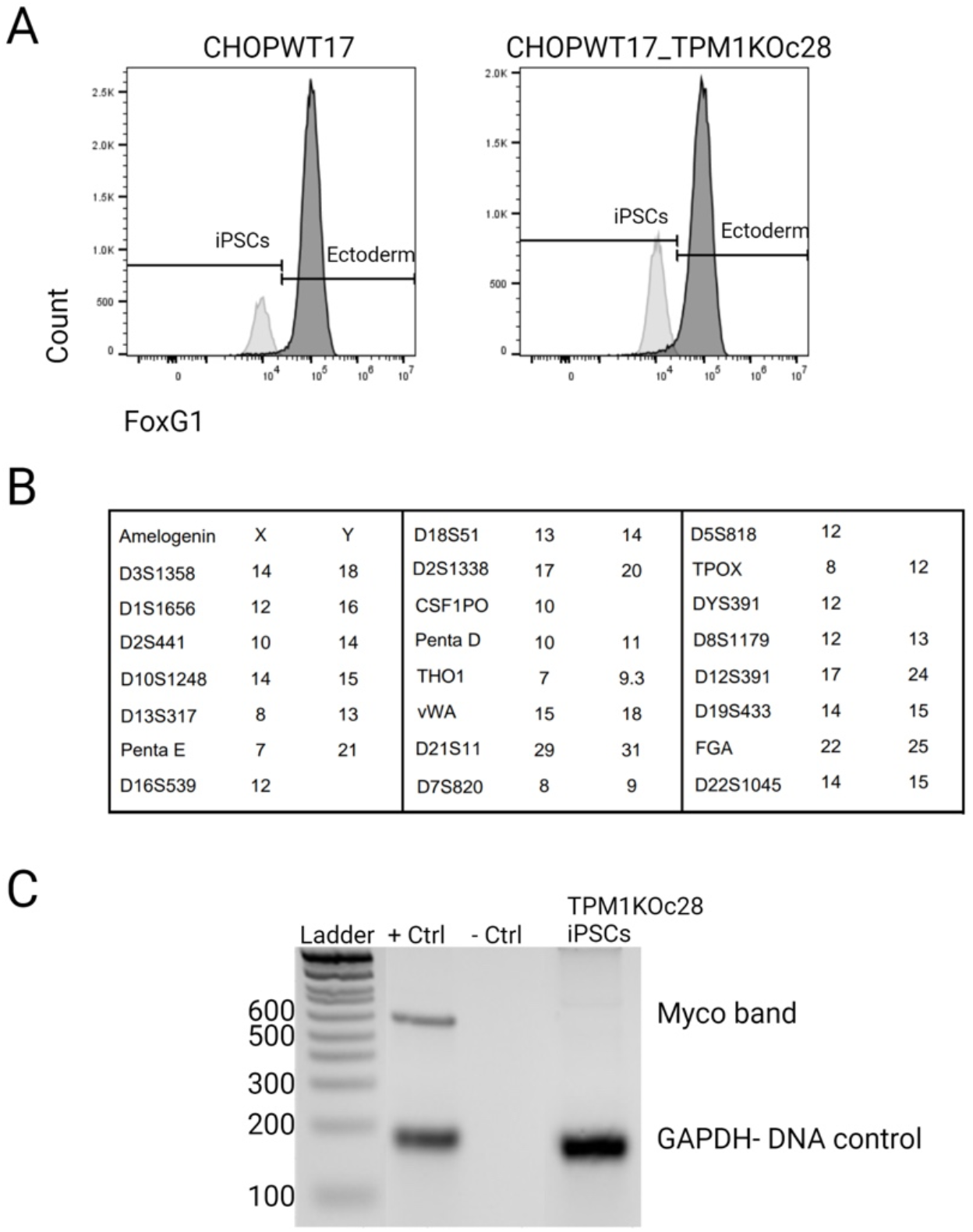
A. Directed differentiation shows efficient production of FoxG1+ ectoderm. B. STR analysis results for CHOPWT17_TPM1KOc28 matched the parental control line. C. Mycoplasma testing was negative for CHOPWT17_TPM1KOc28 iPSCs.

